# Multiple methods for assessing learning and memory in *Drosophila melanogaster* demonstrates the highly complex, context-dependent genetic underpinnings of cognitive traits

**DOI:** 10.1101/2025.02.26.640179

**Authors:** Victoria Hamlin, Huda Ansaf, Reiley Heffern, Patricka A. Williams-Simon, Elizabeth G. King

## Abstract

Learning and memory are fundamental for an individual to be able to respond to changing stimuli in their environment. Between individuals we see variation in their ability to perform learning and memory tasks, however, it is still largely unknown what genetic factors may impact this variability. To gain better insight to the genetic components impacting variation in learning and memory, we use recombinant inbred lines (RILs) from the *Drosophila* synthetic population resource (DSPR), a multiparent mapping population exhibiting natural variation in many traits. Using a reward based associative learning and memory assay, we trained flies to associate an odor with a sucrose reward under starvation condition and measured olfactory learning and memory ability in y-mazes for 50 DSPR RILs. While we do not find significant QTLs for olfactory learning or memory, we found suggestive regions that may be contributing to variability in performance when trained to different odors. We provide evidence that performance with specific odors should be considered different phenotypes and introduce new methods for analysis for olfactory y-maze assays with multiple decision points. Additionally, we compare our data to previously collected place learning and memory data to show there is limited correlation in performance outcomes.

## 1 Introduction

Fundamental cognitive functions such as learning and memory are influenced by a complex interplay between environmental influences and genetics. Learning and memory are critical functions that allow for changes in behavior in response to stimuli encountered in the environment. Performance during learning and memory tasks can show a great deal of variation both across species and between individuals of the same species (Mery 2013; Smid & Vet 2016). Previous behavioral studies using organisms ranging from fruit flies to humans have shown that there is a genetic basis for variation in learning and memory performance, though it is also clear from these studies that the environmental context is also critically important (Burdick *et al*. 2006; Papassotiropoulos *et al*. 2009; Mery 2013; Smid & Vet 2016; Anreiter *et al*. 2017; Williams-Simon *et al*. 2019; Deary *et al*. 2022). To further support the role of a genetic basis, experiments using directional selection have been able to evolve populations to have higher or lower learning performance on very specific tasks (Mery & Kawecki 2002, 2004; Mery *et al*. 2007b; Versace & Reisenberger 2015; Smid & Vet 2016). While there is strong evidence for a genetic basis of phenotypic variation in learning and memory, we still have a limited scope of understanding the suite of specific genomic variants contributing to phenotypic output, due to the highly complex nature of these traits.

Much of our current understanding of the genes impacting learning and memory come from mutational studies where a single gene mutation has a significant impact on learning or memory phenotypes (Davis 2023). These studies provide crucial information on the molecular processes needed for proper learning and memory function, but it is still unclear if allelic variants of these genes contribute to the natural range of phenotypic variation between individuals. As genomic technologies advance, studies have begun to identify potential polymorphisms contributing to variation in learning and memory phenotypes and some mechanisms have been identified. A well-studied polymorphism in the *foraging* gene (*for)* has been shown to regulate differences in multiple behavioral phenotypes across species including *Drosophila melanogaster* (Pereira & Sokolowski 1993; Osborne *et al*. 1997; Anreiter & Sokolowski 2019)*, Mus musculus*(Anreiter & Sokolowski 2019; Duraffourd *et al*. 2019)*, and Homo sapiens* (Sokolowski *et al*. 2017; Anreiter & Sokolowski 2019; Struk *et al*. 2019). Differences in learning and memory have been identified in *Drosophila melanogaster* expressing different *for* alleles; flies with the *for^R^* allele show higher short-term memory performance, increased learning indexes on olfactory appetitive tasks, indifference to social learning, and decreased learning and memory with repetitive training while flies with the *for^s^* allele have higher long-term memory performance, increased learning and memory with repetitive training, and increased performance in social learning (Osborne *et al*. 1997; Kaun *et al*. 2007; Mery *et al*. 2007a; Reaume *et al*. 2011; Kohn *et al*. 2013; Anreiter & Sokolowski 2019; Struk *et al*. 2019). Two intronic polymorphisms in the human dopamine receptor DRD2 have been shown to produce different ratios between the DRD2S and DRD2L splice variants and has been linked to differences in performance on cognitive and attention related tasks (Zhang *et al*. 2007). Continuing to investigate naturally occurring allelic variants and polymorphisms will be critical to understanding the phenotypic variation between individuals in populations.

To identify genomic regions contributing to variation in phenotypes, genome wide association studies (GWAS) and quantitative trait loci (QTL) mapping are methods using natural populations to narrow in on potential causative regions and/or polymorphisms. These methods have been used to study various neurological phenotypes including social behavior (Ruiz-Opazo & Tonkiss 2006; Knoll *et al*. 2018), spatial learning and navigation (Milhaud *et al*. 2002; Ruiz-Opazo & Tonkiss 2006), risk for neurodegenerative and neuropsychiatric disorder (Ramanan & Saykin 2013; Wightman *et al*. 2021; Yao *et al*. 2021), and general cognitive function (Davies *et al*. 2015, 2018). Previous work from our lab identified several QTLs that influence overall performance in a heat box spatial learning and memory test which utilizes an aversive conditioning paradigm (Williams-Simon *et al*. 2019). This study found 16 QTLs that contributed to either learning or memory with five of these QTLs contributing to both phenotypes. In this study, we aim to identify QTLs contributing to olfactory learning and memory phenotypes and compare between the two datasets allowing for identification of overlapping QTLs which may indicate core components leading to phenotypic variation that is shared between multiple learning and memory mechanisms.

To detect QTLs for learning and memory phenotypes, there are several conditioning paradigms that can be used with adult flies that allow for large sample sizes to be obtained (Waddell & Quinn 2001; Pitman *et al*. 2009; Kottler & van Swinderen 2014). Each of these associative learning mechanisms utilize different sensory pathways to intake information from the environment and perception of this information is influenced by pairing of a positive or negative stimulus (Pitman *et al*. 2009; Li & Liberles 2015). While there will be distinct genetic and neuronal pathways activated during these associative events dependent upon the sensory system engaged, there is evidence of overlapping mechanisms central to processing (van Swinderen *et al*. 2009; Aso *et al*. 2014b; Liefting *et al*. 2018). In insects, the mushroom bodies have been shown to play a central role in associative learning by integrating sensory information, assigning valance to cues, and transmitting signals that modulate behavioral outcomes (Aso *et al*. 2014b, 2014a; Vogt *et al*. 2014). Additionally, biogenic amines, such as dopamine, has evidence of roles in both aversive and appetitive memory formation (Unoki *et al*. 2005; Kim *et al*. 2007; Bromberg-Martin *et al*. 2010). This evidence of overlap leads us to hypothesize there will be some shared genetic variants influencing overall variability in general learning and memory phenotypes.

In this paper, we investigate olfactory learning and memory phenotypes in the DSPR lines with the aim of identifying genetic loci contributing to these phenotypes through QTL mapping. Additionally, we aim to compare the olfactory QTL data with previous place learning and memory QTL data collected in the lab to identify any overlapping loci that may contribute to broader learning and memory mechanisms.

## 2 Methods

### 2.1 Mapping Population

We used Recombinant Inbred Lines (RILs) from *population A* of the *Drosophila* Synthetic Population Resource (DSPR) (King *et al*. 2012b, 2012a; Williams-Simon *et al*. 2019; “DSPR” 2023). These RILs were developed through an eight-way intercross which used eight inbred founder lines from locations around the world (King *et al*. 2012b). The founder lines were crossed and randomly mated for 50 generations followed by 25 generations of inbreeding (King *et al*. 2012b; Williams-Simon *et al*. 2019). The founder lines are fully sequenced and the RILs genotyped at over 10,000 SNPs which is used to assign the most likely founder ancestry at all genomic locations for each RIL using a hidden Markov model developed by King *et al*. (King *et al*. 2012a). In the place learning and memory testing, 741 RILs from population A were used. We then selected 50 of these RILs for the olfactory learning and memory testing based on the RILs performance in the place testing and predicted founder ancestry at four previously identified quantitative trait loci (QTLs). To optimize our line choice, we used the following steps: [1] obtained peak information for the QTLs of interest, [2] identified the haplotype ID for each RIL at the peak position, [3] computed the empirical cumulative distribution percentile for both phenotypes using the *ecdf* function from the stats (R Core Team 2022) package then took the mean to create a cumulative score for each RIL, [4] subsetted the RILs based on haplotype ID at the QTLs of interested into a high and low scoring group from the cumulative score, [5] filtered out RILs that had mixed haplotypes, meaning at one QTL they had a haplotype of a high performer while at another QTL they had a haplotype of a low performer to create a list of our top and bottom 25 RILs.

### 2.2 Fly Husbandry/Maintenance

Flies for the olfactory testing were raised on a standard cornmeal yeast diet and kept at 23°^C^ and a minimum humidity of 50% in small vials. Sixteen days psrior to the start of the behavioral assay, 6 males and 10 females were transferred into a new vial and allowed to lay eggs for seven days, then removed to ensure only the F1 generation remained in the vial. Both male and female flies were assayed together and range from 16-22 days post oviposition.

### 2.3 Phenotyping Olfactory Learning & Memory

#### 2.3.1 Conditioning Assay

We used an olfactory appetitive conditioning behavioral assay (Pitman *et al*. 2009; Krashes & Waddell 2011) to condition flies to associate specific odors by pairing with a sucrose reward (Figure 1A). In previous studies, odors were balanced prior to each test by altering concentrations of the odor until the flies divide equally between the two odor arms of the testing apparatus (Pitman *et al*. 2009; Krashes & Waddell 2011; Mohandasan *et al*. 2022). Because of the scale of our study and number of different genotypes being tested, we performed odor balancing on a synthetic population of flies to determine a standard concentration to test all genotypes. In our study we tested each RIL on a standardized concentration of 0.06% OCT and 0.09% MCH diluted in mineral oil.

**Figure 1.**
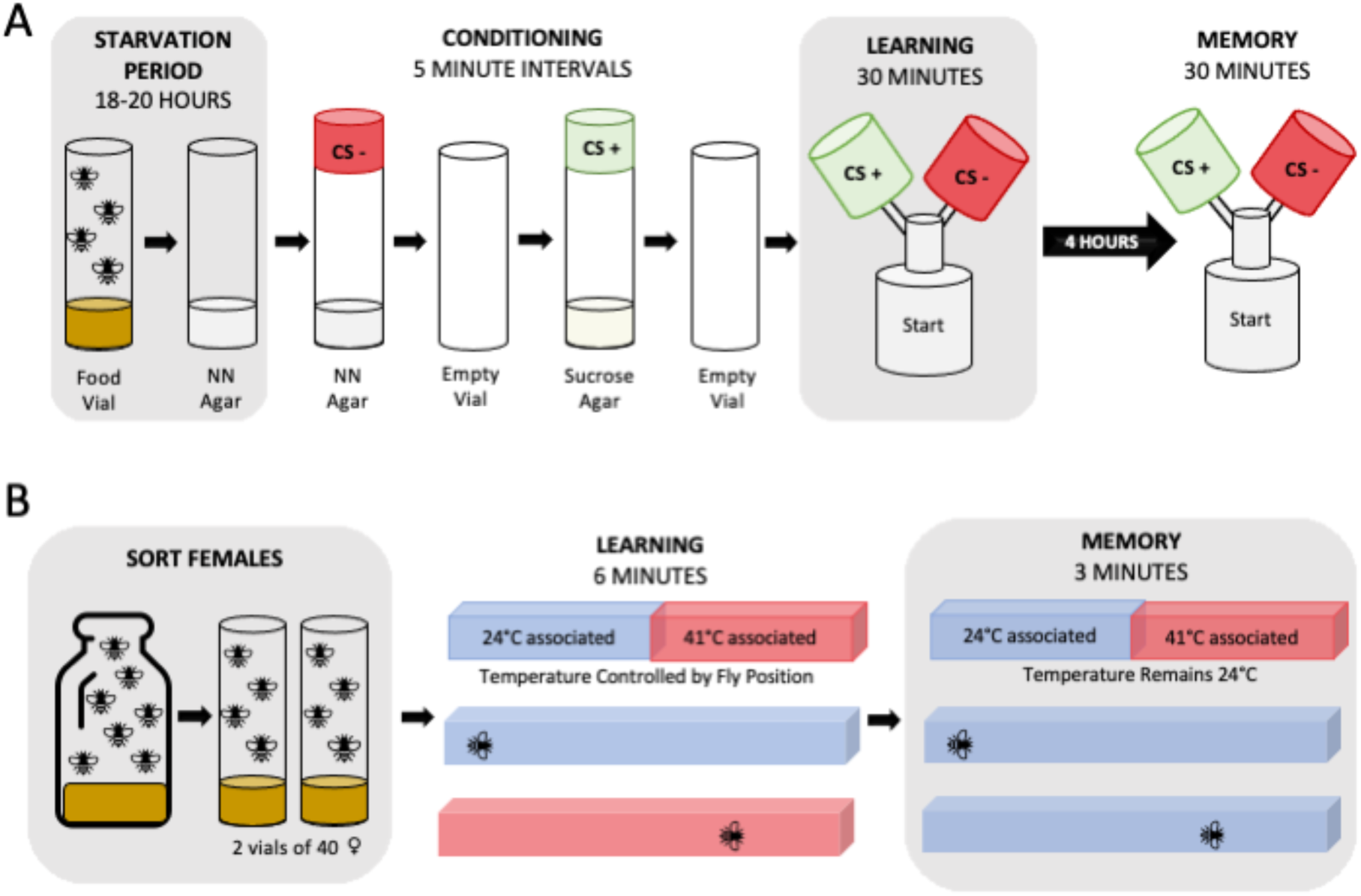
Schematic of the two phenotyping processes. **(A)** Olfactory phenotyping. 60-80 flies were transferred to non-nutritional agar (NN Agar) 15 to 19 days post oviposition for an 18-to-20-hour starvation period. Flies undergo conditioning with each step taking 5 minutes. Conditioning starts with exposure to the negative conditioned stimulus (CS-) odor in NN agar vials, rest in an empty vial, exposure to the positive conditioned stimulus (CS+) in sucrose agar vials, and then a final transfer to an empty vial. The learning assay conducted immediately after conditioning allows flies 30 minutes to choose between the CS+ and CS-odors in the upper chambers of the y-maze. There is a 4-hour rest period before flies undergo the memory assay where they again have 30 minutes to choose between the CS+ and CS-odors. **(B)** Place phenotyping. Approximately 80 female flies are sorted into two food vials 14 days post oviposition. Flies were phenotyped individually in the heat box for 9.5 minutes. In the learning assay, when a fly resides on the cool associated (blue) side of the chamber the box with be 24°C or 41°C when on the hot associated (red) side of the chamber. In the memory assay, the chamber will remain 24°C regardless of fly position.

Flies were transferred to a non-nutritional agar 18-20 hours prior to the start of conditioning. To prepare the conditioning assay, vials of 2M sucrose agar and additional non-nutritional agar vials were placed into fly cages overnight for “pre-flying”. On the day of conditioning, odor caps containing diluted 3-Octanol (OCT) (Millipore-Sigma 8.21859) or 4-Methylcyclohexanol (MCH) (Sigma-Aldrich 153095) were placed on the vials and allowed 20 minutes to odorize. Flies underwent four sequential phases during conditioning protocol: [1] flies are placed into the negative condition stimulus (CS-) vial with non-nutritional agar for five minutes, [2] flies are transferred to an empty vial with fresh air for five minutes, [3] flies are placed into the positive condition stimulus (CS+) vial for five minutes with 2M sucrose agar, and [4] flies are transferred to an empty vial with fresh air for a minimum of 5 minutes before transfer into the testing phase. Each RIL was divided into three groups – two were reciprocally trained to associate either OCT or MCH to the sucrose reward and the third group was unconditioned.

#### 2.3.2 Y-maze Assays

Assays to test learning, memory, and baseline odor preference were conducted in 3D printed Y-mazes (Mohandasan *et al*. 2020, 2022) (Figure 1A) which has two odor chambers in the top arms and the starting chamber at the base. All Y-mazes were odorized for 20 to 30 minutes prior to the start of the assay by pipetting 20uL of diluted odorant onto filter paper and placed in the associated odor chamber. The learning test was conducted within 10 minutes of finishing the conditioning assay. Flies were flipped from the final vial of conditioning into the starting chamber at the base of the Y-maze, placed into a slot under a fabric lightbox chamber, then allowed 30 minutes to select between the two odors presented during the conditioning assay. After the 30-minute learning assay concluded, all flies were collected and categorized based on their choice of either CS+, CS-, or no choice (NC) if they remained in the starting chamber. Flies who selected the CS+ chamber were then tested for memory recall 4 hours after the completion of the conditioning assay following the same Y-maze set up as described earlier. Flies were again collected based on their choice of CS+, CS-, or NC.

#### 2.3.3 Scoring Olfactory Phenotypes

To score performance in the y-maze assays, we counted the total number of flies in each of the three chambers at the end of the 30-minute choice period. In the baseline odor preference assay, the remaining flies in the starting chamber we labelled as *NC* flies and the two odor arms were recorded based on the odor of the chamber, *OCT* or *MCH*. With these counts, we calculated a *climbing score*, which is the proportion of flies who moved into either of the odor chambers out of the total number of flies in the assay, and a *preference score* for each odor, which is the proportion of flies choosing one odor out of the total number of flies who moved into either of the odor chambers. In the learning and memory assays, the flies remaining the starting chamber we also labelled as *NC* flies, and the two odor arms were labelled as either *CS+*, the sucrose paired odor, or *CS-,* the non-nutritional agar paired odor. A *climbing score* was calculated in the same manner as the baseline odor preference assay. The *performance score* was calculated for the learning assay and memory assay by determining the proportion of flies choosing the CS+ odor out of the total number of flies who moved into either of the odor chambers. We developed an additional scoring method that allowed us to assign a score to individual flies even though they are tested as a group. In this binomial method flies were individually scored on two decision points. The first scoring point is assessing the choice to move into an odor chamber; flies that remains in the starting chamber have a score of 0 while flies that move into either of the odor chambers, regardless of it trained odor, have a score of 1. The second scoring point is assessing the correctness of the choice; flies that choose the non-nutritional paired odor have a score of 0 while flies that choose the sucrose paired odor have a score of 1.

#### 2.3.4 Preference-Performance Differentials

As previously mentioned, our study tests flies on a standard odor concentration of OCT and MCH rather than balancing odors for each set of flies before each Y-maze run. To evaluate the effect of odor preference on performance, we calculated a preference-performance differential (PPD) by taking the difference in preference score from the performance score. With this calculation, a PPD value closer to zero means flies are selecting closer to their baseline preference, giving us less confidence they are performing based on their training. PPD values further from zero means that the preference and performance scores have a bigger difference allowing us to be more confident flies are not selecting on their baseline preference.

### 2.4 Olfactory Covariate Testing

#### 2.4.1 Sugar Acuity Assay

We made split agar petri dishes containing half non-nutritional agar and half 2M sucrose agar to determine each RILs preference of sucrose. Flies were starved for 18-20 hours then lightly anesthetized with CO_2_ for transfer to the split agar dish for testing. When most flies had recovered, they were given 5 minutes to explore the plate before an overhead picture was taken for scoring later. Pictures were counted to determine how many flies were on the sucrose side, non-nutritional agar side, and a censored count. Censored count included flies who were not in contact with either agar (side or lid of dish), flies that had not recovered from anesthetic, or flies that were on the boundary of where the agars met. A proportion was calculated for the number of flies on the sucrose plate from the total number of uncensored flies.

#### 2.4.2 Odor Acuity Assay

Using the y-maze, each RIL was tested for odor acuity by placing unconditioned flies into the starting chamber of a pre-odorized y-maze with one upper chamber containing either OCT or MCH and the second upper chamber without odorant. Flies were given 30 minutes to choose between the two upper chambers and collected based on their choice of odor-free chamber, odor chamber, or no choice for flies that remained in the starting chamber. Proportion scores for both climbing and chamber selection were calculated. The climbing proportion is calculated from the number of flies entering any upper chamber from the total number of flies starting the assay. The chamber selection proportion is calculated from the number of flies entering the odor chamber from the total number of flies who entered either chamber.

#### 2.4.3 Lightbox Calibration

To eliminate chamber side bias from phototaxis, we performed light box calibration. Unconditioned flies were placed into to the starting chambers of clean y-mazes without odorants in the upper chambers. The y-maze was placed into a slot under the fabric light box and allowed to choose between the upper chambers for 30 minutes. Flies were collected based on the chamber they ended in.

### 2.5 Phenotyping Place Learning & Memory

Place Learning & Memory Phenotype data was previously published and full details can be found in Williams-Simon et al, 2019 (Williams-Simon *et al*. 2019). Flies were trained in individual chambers of a heat box apparatus (Williams-Simon *et al*. 2019) to evaluate place learning and memory (Figure 1B). For this behavioral assay, individual flies are placed into a chamber for a total of 9.5 minutes of testing which has three phases: [1] 30-second pretesting phase, [2] 6-minute learning phase, and [3] 3-minute memory phase. Using infrared light to track the flies’ position, the whole chamber temperature will increase to 41°^C^ when the fly is on the hot associated side then return to 24°^C^ when the fly is in the cool associated side during the learning phase. In the memory phase, the fly’s position was tracked, however, the chamber remained 24°^C^ regardless of its position. Performance Indices (PI) were calculated for learning and memory based off the amount of time a fly spent on each temperature associated side of the chamber during that respective phase. To calculate the PI the time spent on the hot associated side is subtracted from the time spent on the cool associated side divided by total time in the chamber. The maximum score of 1 indicates a total avoidance of the hot associated side while the minimum score of -1 indicates a total preference for the of the hot associated side.

### 2.6 Mapping for Olfactory QTLs

Using the binomial scores for individuals tested from the 50 RILs, we fit two generalized linear mixed models using the glmmTMB package (Brooks *et al*. 2017) at over 11,000 positions across the genome for learning, memory, and climbing phenotypes. The two models we fit were as follows:

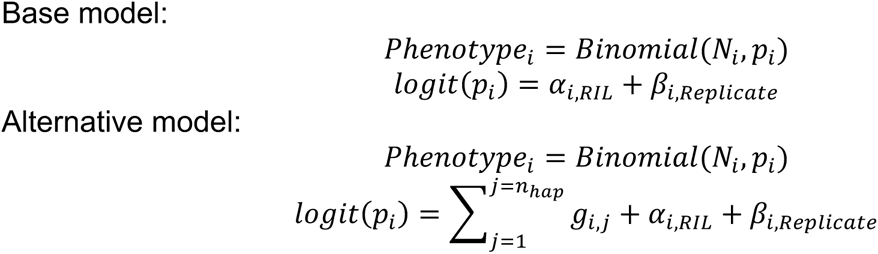

where *i* is the *ith* individual, *RIL* is each DSPR RIL, *Replicate* is each vial replicate, *g_i,j_* is the probability the RIL harbors the *jth* haplotype at the position. Given our sample size, often not all 8 haplotypes are represented at a given position. Thus, we dropped the *g_i,j_* term when fewer than 4 RILs harbored that haplotype at the position to avoid overfitting. Thus, the number of haplotype probabilities included in the model differs across the genome scan and is denoted by *n_hap_*.

Since there is some indication that response to odors may influence learning and memory, we analyzed each odor and test type combination: OCT Learning, MCH Learning, OCT Memory, and MCH Memory. For each of these phenotypes, our base model does not consider haplotype at each position while the second model does. In the second generalized linear mixed model, we test if the probable haplotype at each position can predict the outcome of interest with RIL and Replicate set as random effects in the analysis. From these two models, we obtained the log likelihoods and performed the Likelihood Ratio Test. To perform a QTL scan, we then calculated the negative log_10_ p-value via a chi-square test. To determine the overall significance threshold, we estimated the positive false discovery rate (pFDR) using the q-value package (Storey JD, Bass AJ, Dabney A, Robinson D 2024). To define suggestive QTLs specifically for the difference in performance between odor trainings, we took the absolute value of the difference between the odors negative log_10_ p values at each position. We then selected the top three unique positions for learning and memory that had a difference greater than 2 and analyzed haplotype performance at these positions.

### 2.7 Data Availability

Raw *olfactory* learning and memory phenotypic data are available from zenodo (https://zenodo.org/): http://doi.org.10.5281/zenodo.14907900. Raw *place* learning and memory phenotypic data are available from zenodo (https://zenodo.org/): http://doi.org/10.5281/zenodo.2595557. Founder genotype assignments from the hidden Markov model are available as two data packages in R (http://FlyRILs. org/) and are available from the Dryad Digital Repository (http://dx.doi.org/10.5061/dryad.r5v40). See King et *al.* (King *et al*. 2012b, 2012a) for details of the DSPR datasets. All DSPR data is also available centrally at http://FlyRILs.org. The complete code used to perform all analyses is available at GitHub (https://github.com/EGKingLab/Olfactory_LearnMem).

## 3 Results

### 3.1 General Learning and Memory Patterns

We measured olfactory learning and memory performance and subsequently climbing performance for each test in 50 DSPR RILs and see variability between lines in all phenotypes (Figure 2). In the learning assay, we measured performance of 17,251 flies and our sample size ranged from 64 to 587 flies with an average of 345 flies starting the assay. Learning scores vary between RILs with the lowest performing line scoring 0.399, the highest performing line scoring 0.682, and mean learning performance score of 0.529 on a scale of zero to one. Additionally, the climbing sample size in the learning assay ranged from 11 to 326 flies with an average of 167 flies entering an odor chamber during the assay and the mean climbing performance score of 0.495 (range 0.073 to 0.900).

**Figure 2.**
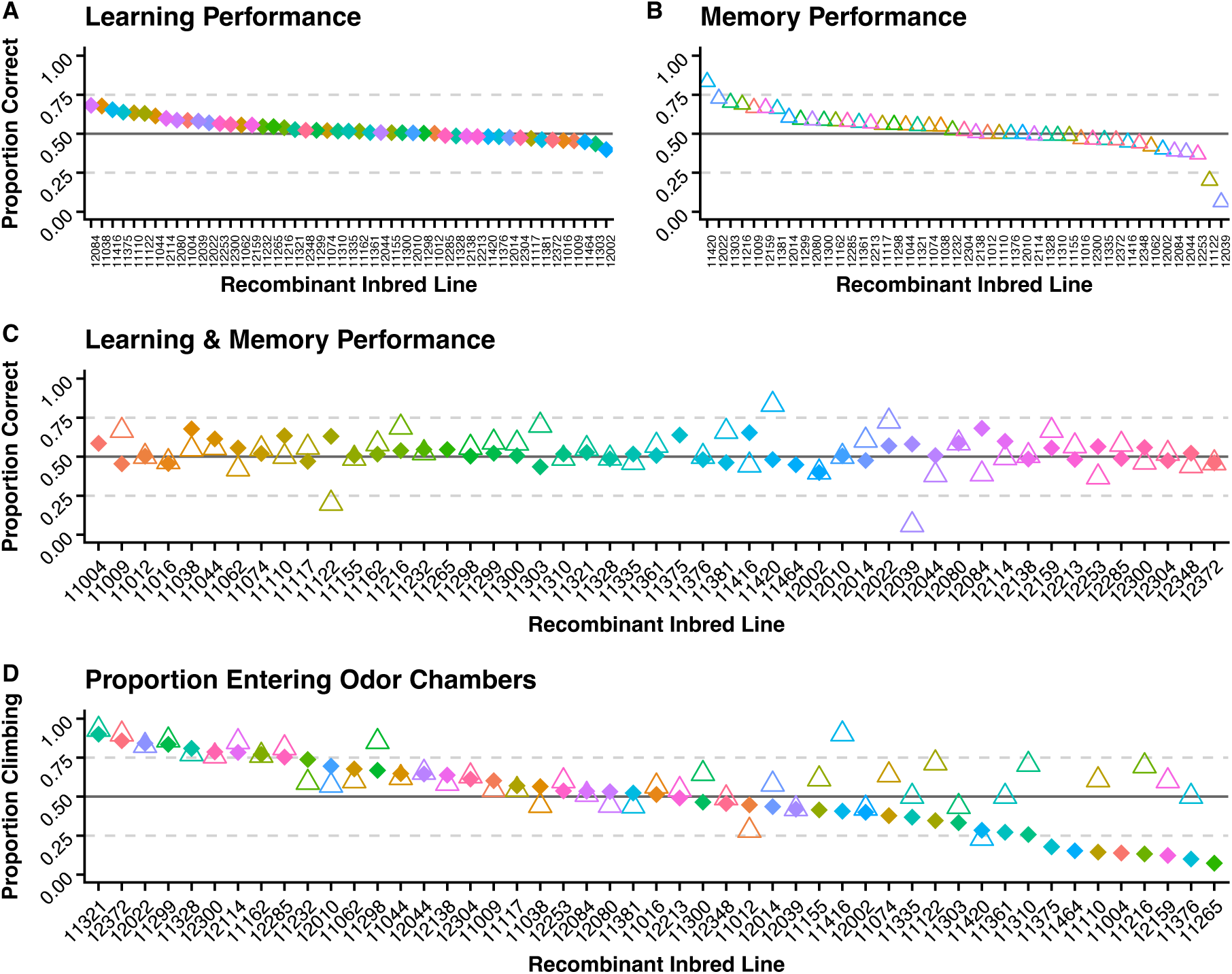
Overall olfactory learning and memory performance. Diamonds represent data collected from learning assays. Triangles represent data collected from memory assays. **(A)** Learning performance scores for each RIL calculated by taking the proportion of flies choosing the sucrose trained odor (CS+) from the total number of flies entering an odor chamber. **(B)** Memory performance scores for each RIL calculated by taking the proportion of flies choosing the sucrose trained odor (CS+) from the total number of flies entering an odor chamber. **(C)** Learning and memory performance scores are plotted together to show correlations between performance on each test. **(D)** The climbing score is the proportion of flies moving into either odor chamber out of the total number of flies starting the test.

In the memory assay, we measured performance of 4,046 flies and our sample size ranged from 10 to 178 flies with an average of 88 flies starting the assay. Memory scores vary between RILs with the lowest performing line scoring 0.063, highest performing line scoring 0.833, and the mean memory performance score of 0.520 on a scale of zero to one. The climbing sample size in the memory assay ranged from 6 to 158 with an average of 55 flies entering an odor chamber and the mean climbing performance of 0.615 (range 0.230 to 0.930).

### 3.2 Baselines for Odor Preference Differ Between RILs

Untrained baseline odor preferences to 3-Octanol (OCT) and 4-Methylcyclohexanol (MCH) were obtained for each of the RILs to evaluate the impact of natural preference on learning and memory performance (Figure 3A). When naïve flies are presented the choice between OCT and MCH, 22 out of 49 RIL fall within a neutral score of 0.45 to 0.55 (0.5 _+/- 10%_). The range for baseline odor preference is 0.184 to 0.713 in which a score of 0 indicates full preference for MCH and a score of 1 indicates full preference for OCT.

**Figure 3.**
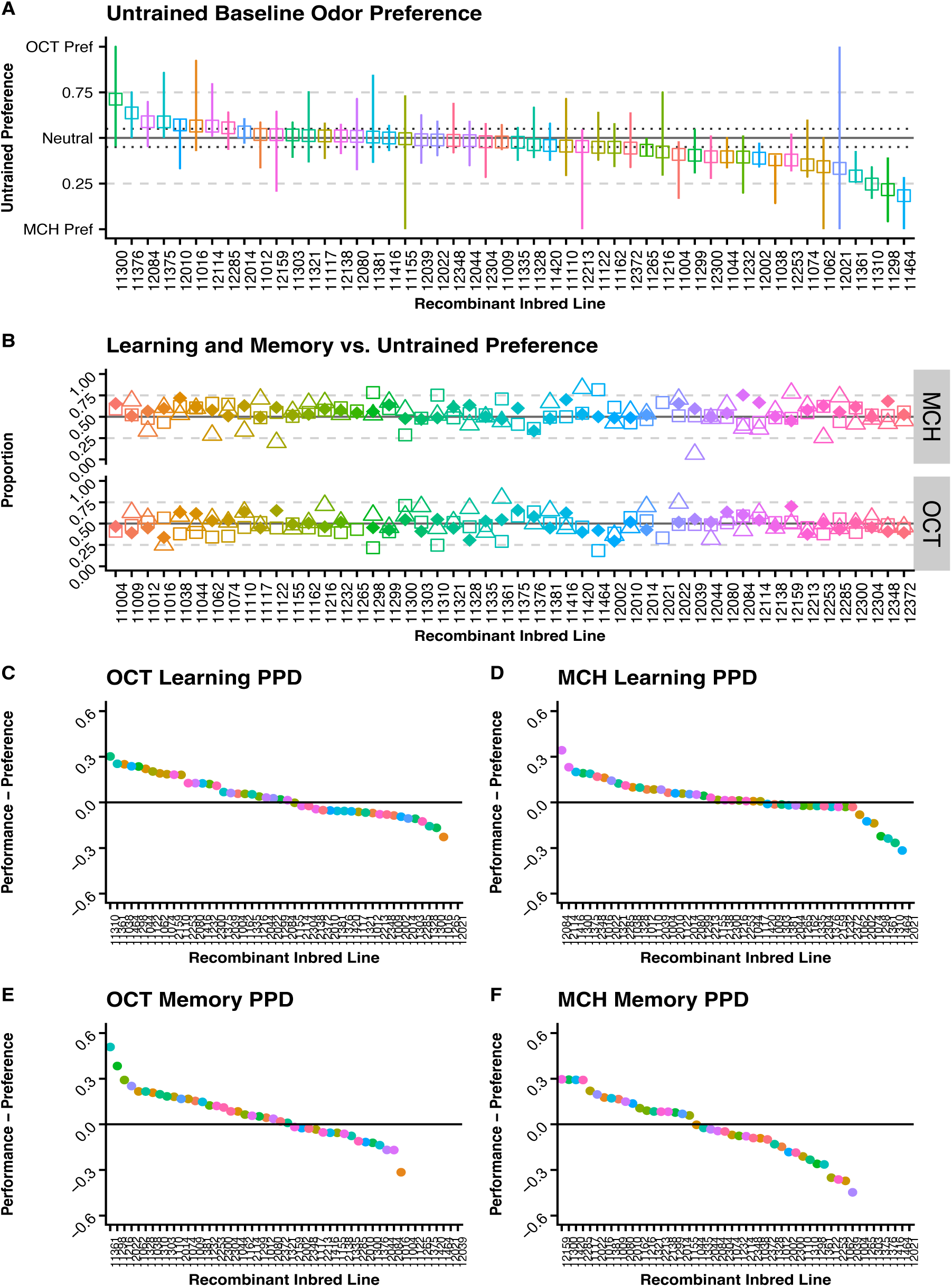
Untrained baseline odor preference impact on learning and memory. Diamonds represent data collected from learning assays. **(A)** Untrained odor preference for each RIL. Squares indicate cumulative score for the RIL and lines represent the range of scores across multiple replicates. **(B)** Untrained odor preference(squares) plotted with learning(diamonds) and memory(triangles) performance. **(C-F)** Preference Performance Differentials for each odor and testing pair.

In Figure 3B, preferences to the trained odor are plotted with the square while learning and memory scores are plotted with the diamond and triangle, respectively. This visualization shows us how closely the learning and memory scores are to the baseline odor preference. When the baseline odor preference is distant from the learning and memory scores this provides some indication that flies are responding to the conditioning paradigm, however, when these preferences align closely with learning and memory performance we cannot confidently determine if they are responding to training or just selecting based off natural preferences. Our data shows that several of the lines have learning and memory scores that are closely aligned with their baseline odor preferences, therefore we explore further methods of analysis to delineate lines that are strong performers when trained against their natural preference.

Two methods of analysis were used in evaluating the impact of untrained baseline odor preference on the performance in olfactory learning and memory assays. First, we calculated a Performance Preference Differential (PPD) shown in Figure 3C-F to evaluate how different the RILs performance score was compared to their preference score. Scores father from zero indicate lines with bigger differences between preference and performance scores giving more confidence that they are not relying only on baseline preferences in the test. In the second method used to evaluate the impact of untrained baseline odor preference on the performance, we looked for RILs with a performance proportion above 0.5 and a preference proportion under 0.5 – meaning this group selected the trained sucrose paired odor even though their baseline preference was to the reciprocal odor (Figure 4A-D). This data suggest these flies are stronger performers in that test because they can “override” their preference and select the sucrose paired odor. In Table 1, the list of RILs that meet this requirement is listed for each preference and test type. For flies who had a preference to MCH but performed well when trained to OCT 7 RILs were identified for performing well in the learning assay, 5 RILs were identified for performing well in the memory assay, and 9 RILs performed well on both assays. For flies who had a preference to OCT but performed well when trained to MCH 4 RILs were identified for performing well in the learning assay, 5 RILs were identified for performing well in the memory assay, and 6 RILs performed well on both assays.

**Figure 4.**
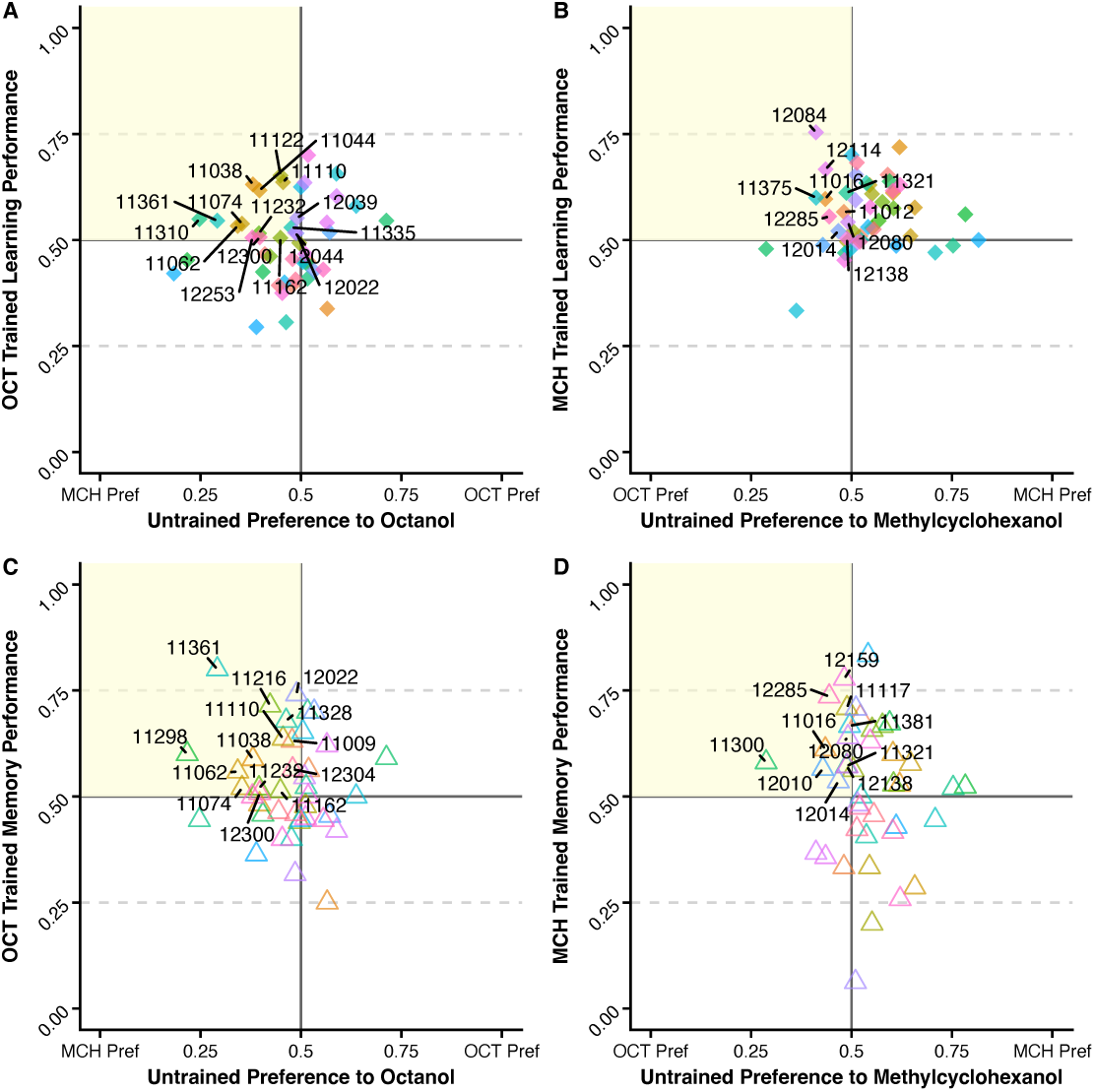
Identifying RILs with good performance when trained against preferences. Labelled RILs have a performance score above 0.5 and a preference score below 0.5 – indicating the flies are selecting the sucrose trained odor which is opposite to their preferred odor **(A)** Highlighted RILs trend towards MCH preference but chose OCT after sucrose pairing during the learning assay. **(B)** Highlighted RILs trend towards OCT preference but chose MCH after sucrose pairing during the learning assay. **(C)** Highlighted RILs trend towards MCH preference but chose OCT during the memory assay. **(D)** Highlighted RILs trend towards OCT preference but chose MCH during the memory assay.

**Table 1.**
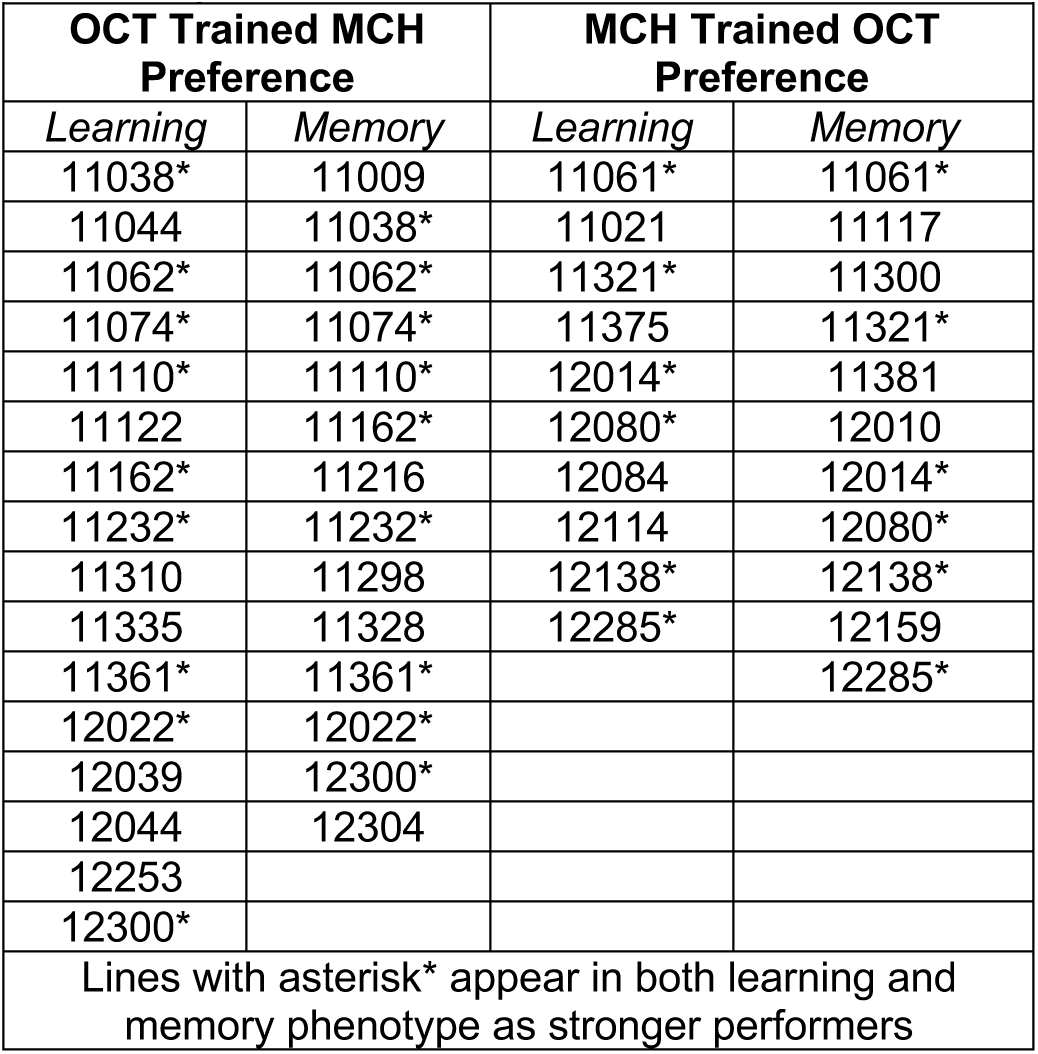
List of RIL with flies who performed well when trained against baseline odor preference.

| **OCT Trained MCH Preference** |  | **MCH Trained OCT Preference** |  |
| --- | --- | --- | --- |
| *Learning* | *Memory* | *Learning* | *Memory* |
| 11038* | 11009 | 11061* | 11061* |
| 11044 | 11038* | 11021 | 11117 |
| 11062* | 11062* | 11321* | 11300 |
| 11074* | 11074* | 11375 | 11321* |
| 11110* | 11110* | 12014* | 11381 |
| 11122 | 11162* | 12080* | 12010 |
| 11162* | 11216 | 12084 | 12014* |
| 11232* | 11232* | 12114 | 12080* |
| 11310 | 11298 | 12138* | 12138* |
| 11335 | 11328 | 12285* | 12159 |
| 11361* | 11361* |  | 12285* |
| 12022* | 12022* |  |  |
| 12039 | 12300* |  |  |
| 12044 | 12304 |  |  |
| 12253 |  |  |  |
| 12300* |  |  |  |
| Lines with asterisk* appear in both learning and memory phenotype as stronger performers |  |  |  |

### 3.3 Covariate Acuity Measures and Correlations

Odor acuity was measured with a y-maze set up with one upper chamber containing the odorant and the second upper chamber without odorant (null chamber). A proportion score was used to measure odor acuity with a score of 1 indicating all counted flies were in the odor chamber upon collection while a score of 0 indicating all counted flies were in the null chamber upon collection. Overall, we see variability between RILs in acuity to both odors used in our assay. For OCT, the range of proportion scores was 0.00 to 0.82 with a mean of 0.44 (Figure 5A). For MCH, the range of proportion scores was 0.26 to 1.00 with a mean of 0.51 (Figure 5A). We plotted odor acuity scores against learning and memory scores to determine if there is correlation between acuity and performance and find there is no linear correlations between odor acuity and learning or memory (Supplemental Figure 5C-D).

**Figure 5.**
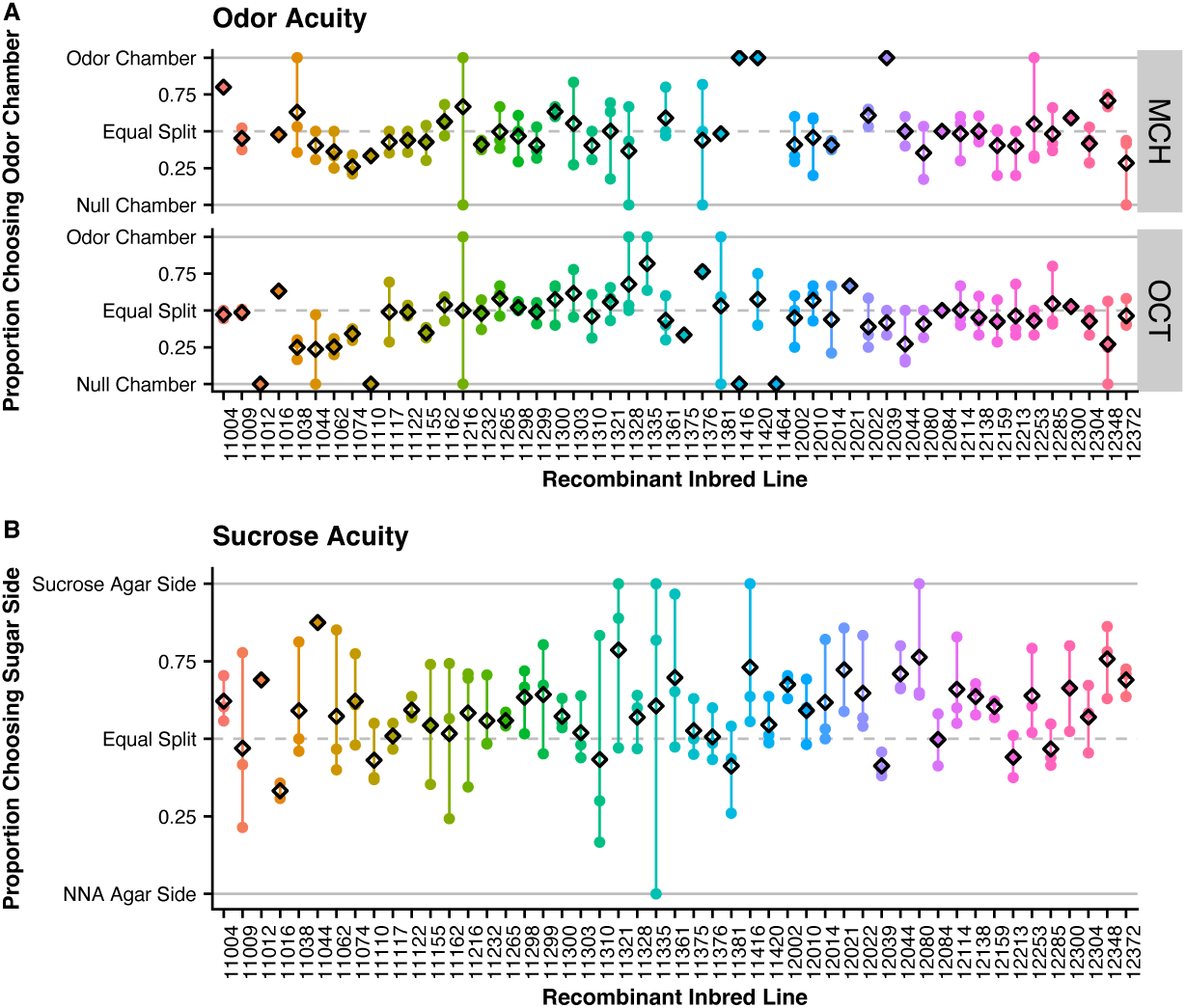
Acuity Plots showing each replicate score with circles colored by RIL and the average score for the replicate indicated by the black diamond. **(A)** Odor acuity for MCH and OCT are variable between RILs. **(B)** Sucrose acuity for starved flies is variable by RIL.

Sucrose acuity was measured by placing starved flies on a split agar plate for 5 minutes. A proportion score was used to measure acuity with a score of 1 indicating all counted flies were on the sucrose side of the plate while a score of 0 indicating all counted flies were on the non-nutritional agar side of the plate. Overall, we see variability in sucrose acuity between RILs with proportion scores ranging from 0.33 to 0.88 and mean of 0.59 (Figure 5B). We plotted sucrose acuity scores against learning and memory scores to determine if there is a correlation between sucrose detection acuity and performance and find there is no linear correlations between sucrose acuity and learning or memory (Supplemental Figure 5E-F).

### 3.4 QTL Mapping & Haplotype Analysis

We performed genome scans using a generalized linear mixed model to analyze the binomial phenotype data collected for learning, memory, and climbing phenotypes and mapped these results for QTLs. Our calculated False Discovery Rate (FDR) was 0.89 and we were unable to detect and any significant QTLs for olfactory learning (Figure 6A), olfactory memory (Figure 6B), or climbing (Supplemental Figure 4) at this threshold. Additionally, because of this high FDR, we did not consider any peaks to be suggestive QTLs for overall olfactory learning, olfactory memory, or climbing. When comparing the QTL maps of performance on learning and memory assays when trained to different odors we see regions where there are differences in p-values that may be indicative of areas involved in modification of behavior dependent on the training odor and/or areas involved in odor preference. Taking the absolute value of the difference between negative log_10_ p-values, we defined three suggestive differential peaks for each phenotype by selecting the top three unique regions with a difference in value above 2. Since these peaks are only suggestive, we further investigate by visualizing each haplotypes mean phenotype score at these suggestive peaks, L1-L3 for learning and M1-M3 for memory (Figure 6). At the L1 and L2 suggestive peaks, we see variability in the mean proportion correct between haplotypes when trained to OCT, however, when trained to MCH the mean proportion correct between haplotypes is approximately equal at these positions. This suggests that these positions may play a role in influencing performance when OCT trained but a more limited role when MCH trained. We see the opposite trend at the M2 position where the mean proportion correct is about equal for OCT trained but variability can be seen between haplotypes when MCH trained. At the M1 and M3 suggestive peaks, we see variability in mean performance between haplotypes for both OCT and MCH meaning this could be an area influencing a more general olfactory memory mechanism. Lastly, L3 shows that both OCT and MCH trained have limited variation between haplotypes so is unlikely to be a region influencing variation in learning performance. All haplotype mean performance visualizations are shown in Supplemental Figure 2.

**Figure 6.**
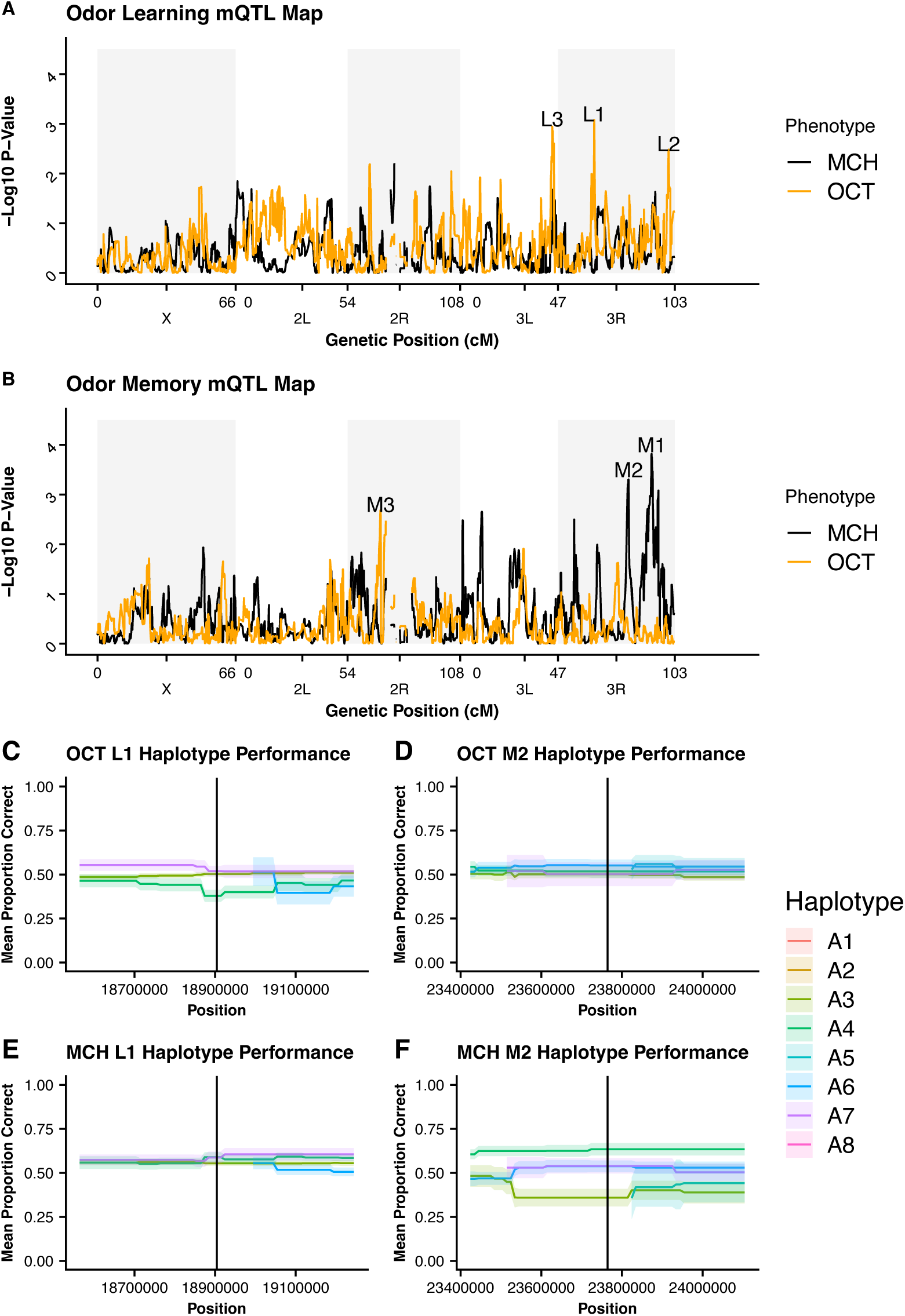
Learning and Memory Performance by Odor **(A)** Learning QTL Map for OCT and MCH trained flies. Odor suggestive peaks L1, L2, and L3 represent the top three positions where p-values differ the most between the two odor trainings. **(B)** Memory QTL Map for OCT and MCH trained flies. Suggestive peaks M1, M2, and M3 represent the top three positions where p-values differ the most between the two odor trainings. **(A,B)** No significant QTLs were identified with pFDR value of 0.89. **(C,E)** Comparison of haplotype performance at L1 suggestive peak. More variability is seen between haplotypes when trained to OCT at this position. **(D,F)** Comparison of haplotype performance at M2 suggestive peak. More variability is seen between haplotypes when trained to MCH at this position.

Previous work in our lab identified several QTL associated with place learning and memory. In Figure 7, we add the place learning and memory data to our olfactory QTL maps to identify if there are any overlaps in genomic regions that may contribute to overall mechanisms involved in place learning and memory. Since there are no significant QTLs found for the olfactory learning and memory data, we cannot confidently determine if there are common regions contributing to overall learning and memory phenotypes. When selecting the RILs used for olfactory learning and memory, we preselected haplotypes at four QTL positions based on prior performance on place learning and memory testing, therefore we used the haplotype mean performance visualization to compare performance index scores across the three phenotypes to see if shared patterns arise at these selected locations (Figure 7, Supplemental Figure 3). At the Q4 learning peak we see place learning and OCT learning have a variant in different haplotypes that drive a difference in performance while MCH haplotypes are performing similarly at this position. At the Q6, Q14 and Q16 peaks we see that place and MCH learning have haplotype variant attributed to difference in performance while haplotypes in OCT are preforming similarly. For the Q4 memory peak we see variants in place learning leading to differences in performance where both olfactory memory phenotypes have similar performance among haplotypes. All QTL haplotype mean performance visualizations are provided in Supplemental Figure 3.

**Figure 7.**
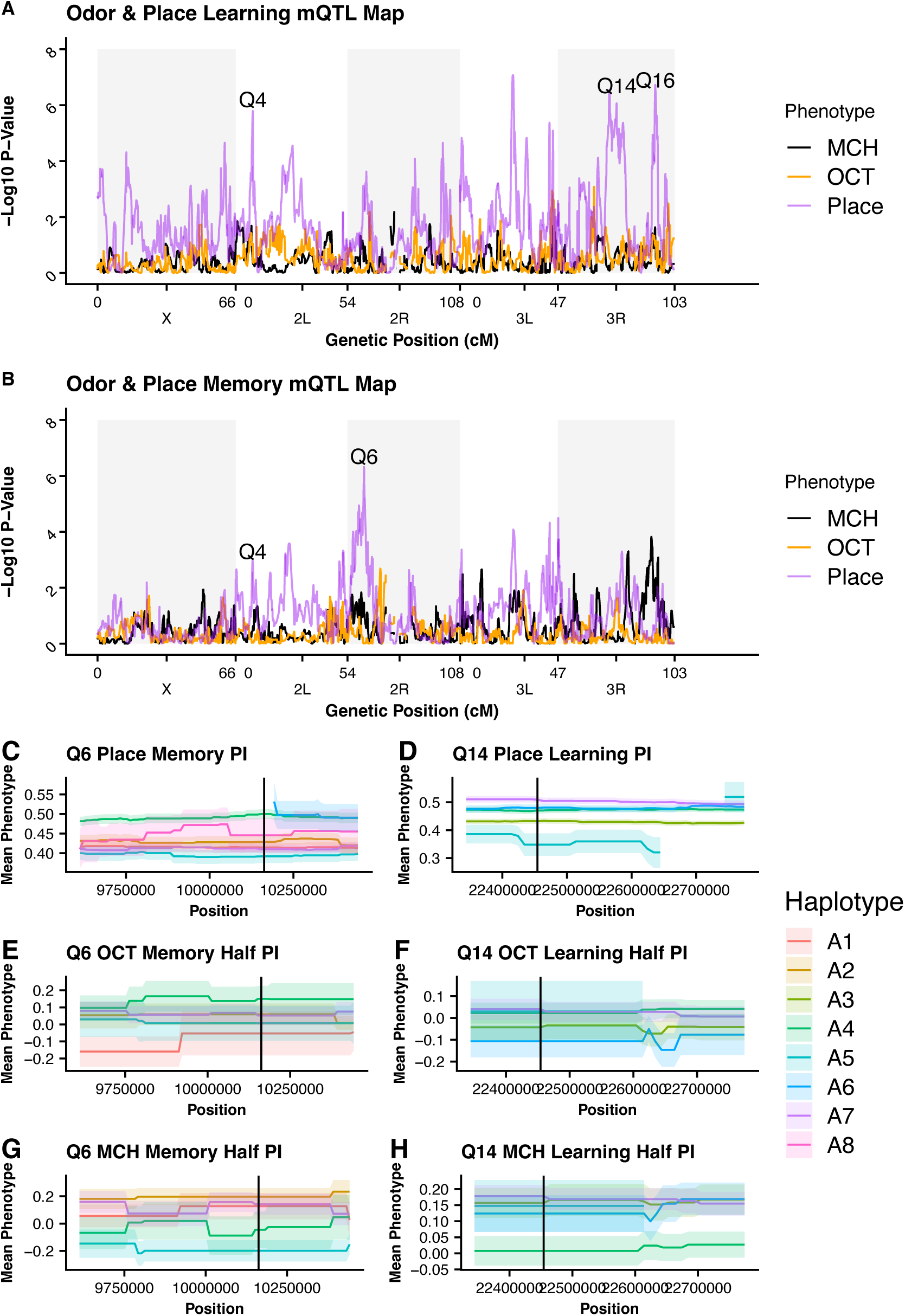
Olfactory and Place Learning and Memory Comparisons. **(A)** QTL map for Place, OCT, and MCH learning. Q4, Q14, and Q16 represent the Place Learning QTLs used to select RILs for olfactory testing. **(B)** QTL map for Place, OCT, and MCH Memory. Q4 and Q6 represent the Place Memory QTLs used to select RILs for olfactory testing. **(C,E,G)** Haplotype comparison at Q6 position for all phenotypes. **(D,F,H)** Haplotype comparison at Q14 position for all phenotypes.

When assessing the QTLs and suggestive peaks with the haplotype performance visualization, regions where we see one or more haplotypes with performance differences beyond the standard error of 1, we presume that there is a variant in that haplotype affecting the performance on the specific assay. When comparing across phenotypes, we see that different haplotypes can be involved in driving performance differences seen at these positions: this could be different variants within the same gene/regulator or from separate components within this peak. It is important to note that this method does not detect down to single nucleotides so the exact variant cannot be defined without further investigation.

## 4 Discussion

Measuring learning and memory phenotypes can be complex due to the many components influencing the organism’s ability to perform the studied behavior such as innate preferences, motivation, and locomotor capabilities. In previous work, the performance index (PI) was used to score learning and memory behaviors, however, we believe this method was missing key components of the phenotype when utilizing Y-mazes for learning and memory experiments. The first issue is that PI will only account for flies that make it into an odor chamber, while this method works well for T-maze assays where flies can only be in one of two chambers, in the Y-maze flies can remain in the starting chamber and therefore are unaccounted in the measurements. This leads to the second issue where the PI method does not handle differences in sample size well; when smaller samples are used the resulting score can become inflated in comparison to larger samples (Supplemental Figure 1). With the y-maze, if we place the same number of flies into a starting chamber, the final number of flies finishing in the odor chambers is highly variable between each replicate and each RIL tested making it impossible to control for the sample size. Lastly, the PI method averages the scores from the two odor trainings to get a final PI, however, performance with a specific trained odor can be influenced by baseline odor preferences. Traditionally, the balancing of odors should reduce this impact, but in our work, we chose to use a standard concentration due to the volume of flies being tested with different genetic backgrounds. Each genotype exhibits differences in untrained odor preference when choosing between OCT and MCH, as seen in Figure 3A, which would require different odor concentrations to be used in a single testing session if using the odor balancing method. Additionally, this can mask differences in performance between the two odors which may remove relevant ecological context.

Due to these pitfalls, we decided to use a binomial scoring method which allows us to account for many of the above-mentioned issues in the following ways: [1] to address the issue of unaccounted flies in the assay, we split our assay into two choice points for analysis, a climbing choice point and a correctness choice point. The climbing choice scores the fly on whether they entered an odor chamber or remained in the starting chamber. This should be considered separately from learning and memory scoring because there are additional factors that can contribute to flies remaining in the starting chamber such as general activity levels, motivation, or exploration behaviors. The correctness choice point scores the fly based on whether it selected the sucrose agar associated odor or the non-nutritional agar associated odor. [2] To address the issue in sample size, the binomial scoring method assigns each individual fly a score even though they have been tested in groups. Since this score is applied to an individual fly, the fluctuation in sample size does not influence the final score the fly receives because it is not a part of the calculation. [3] To address the issue of difference in odor performance, we analyze each RILs performance on odors individually and in relation to its untrained baseline odor preference. Additionally, the binomial method allows for additional analysis of climbing behavior by adding the choice score and role of untrained preferences because of the individual odor analysis giving us deeper understanding of the contribution from these components leading to the overall learning and memory output.

Our study shows that RILs exhibit variability in olfactory learning and memory performance. While we did not find significant quantitative trait loci associated with olfactory learning or memory, when comparing performance between trained odors, there are suggestive peaks indicating some genetic positions may influence learning and memory performance differently based on which odor is paired to the sucrose reward. When looking at the haplotype analysis in the six suggestive regions we see three of our peaks show variability in performance between haplotypes in one odor but not the other and two peaks where both odors have variability in performance between haplotypes (Supplemental Figure 2). We also show that RILs exhibit variability in untrained baseline odor preferences which can influence performance and interpretation of learning and memory assays. When analyzing a RILs performance, it is important to consider its baseline odor preference because a fly scoring well when trained to the odor they naturally prefer can cause a misguided interpretation of that fly being a strong performer where that may not be the case. In Figure 4, we plot the preference vs performance scores to see which RILs have a performance proportion of 0.5 or higher when trained to the odor opposite of their baseline preference; 26 of 49 of RILs for learning (53%) and 15 of 45 RILs (33%) for memory met this criterion. However, when looking at these scores without baseline consideration we see 33 of 49 RILs (67%) have a performance proportion greater 0.5 for learning and 24 of 45 RILs (53%) for memory which means we can be inaccurately categorizing these additional lines as strong performers when they are potentially only acting on their innate preference. This indicates that performance on each trained odor should be considered a different phenotype, baseline preferences should be considered in evaluation of performance, and averaging of scores could be masking important context dependent information necessary to fully understand the implementation of these traits in an ecological setting. Additionally, the variability of performance seen between RILs highlights the importance of including many genotypes in studies trying to understand complex traits given the genetic background plays a significant role in phenotypic presentation. Limiting studies to a single genotype or single baseline can severely hinder our ability to extrapolate general principles guiding development of complex phenotypes such as olfactory learning and memory.

When comparing olfactory learning and memory to place learning and memory we did not see a significant overlap in significant or suggestive odor peaks (Figure 7A-B) and found no linear correlation between performance scores of each phenotype (Supplemental Figure 5A-B). This lack of correlation indicates there is no singular haplotype linked to being the “best” performer on these assays, rather it indicates a diverse array of haplotypes are capable of performing well and these haplotype groups can vary between different testing paradigms. While there is evidence from previous studies of potential shared networks for overall learning and memory (van Swinderen *et al*. 2009; Aso *et al*. 2014b; Liefting *et al*. 2018), reasons for this lack of significant shared peaks could have both biological and/or experimental explanation. Both assays used an associative conditioning paradigm but used different stimuli, aversive for place testing in contrast to appetitive reward for olfactory. Additionally, these assays targeted different sensory systems, thermosensation in place testing and olfactory and gustatory systems in the olfactory testing. Targeting these different sensory systems with contrasting modalities of conditioning require different circuits and receptors for proception and may limit how much overlap we are able to detect. Experimentally, we had preselected for RILs with specific haplotypes at peaks from place learning study which limited the genetic variability at these four place QTLs and only tested 50 RILs in comparison the place work which tested 741 RILs. By only studying a small subset we may have limited our detection power if other haplotypes were stronger contributors to the olfactory phenotype. Through the haplotype analysis at the previously identified place learning and memory QTLs, we were able to identify some shared trends where single haplotypes can be identified as contributors to difference in performance, though it was typically not the same haplotype contributing between place and olfactory phenotypes (Supplemental Figure 3).

Overall, in this study we show that there is highly complex variability influencing olfactory learning and memory performance highlighting the importance of genetic background in phenotypic presentation of complex phenotypes. We have identified five total suggestive regions, two suggestive regions for olfactory learning and three suggestive regions for olfactory memory, through haplotype analysis at peaks with greatest differences between negative log_10_ p-values. Our data indicates each odor should be considered separate phenotypes due to variability in the haplotype performance between odors at a few of the suggestive peaks. Consideration of baseline odor preference in evaluation of olfactory learning and memory phenotypes can be used to better identify if there is response to training verse selection of natural preference.

## Acknowledgements

We thank Dr. Stuart Macdonald for supplying us with the DSPR RILs. We thank Dr. Gaurav Das and Radhika Mohandasan for providing the files for 3D printing Y-mazes and input on setting up Y-maze experiments. This material is based upon work supported by the National Science Foundation under Grant No. IOS-1654866 and by the National Institutes of Health under Grant No. R35GM149238.

