## Supplemental Figures for "Multiple methods for assessing learning and memory in *Drosophila melanogaster* demonstrates the highly complex, context-dependent genetic underpinnings of cognitive traits"

**Supplemental Figure 1**

**PERFORMANCE INDEX CALCULATION**

$$\frac{(Total\ Correct\ - Total\ Incorrect)}{Total\ Number\ of\ Flies\ in\ Odor\ Chambers}$$

**Small Sample Size Effect**

**n=30**

$$a.\ \frac{15-15}{30}=0.000$$

$$b.\ \frac{20-10}{30}=0.333$$

$$c.\ \frac{25-5}{30}=0.667$$

$$d.\ \frac{30-0}{30}=1.000$$

**Large Sample Size Effect**

**n=300**

$$e.\ \frac{150-150}{300}=0.000$$

$$f.\ \frac{155-145}{300}=0.033$$

$$g.\ \frac{160-140}{300}=0.067$$

$$h.\ \frac{200-100}{300}=0.333$$

**Supplemental Figure 1.** Effect on sample size on Performance index. With smaller sample sizes the increase in score can change dramatically with smaller changes in number of correct to incorrect choosers. When comparing RILs which have different samples sizes due to lack of control at this y-maze choice point, RILs with smaller samples sizes can be inflated to seem like they are stronger performers than those with larger sample sizes. While the scores remain the same when the ratios are relative like we see in equations **A&E** and **B&H**, this distortion can lead to misinterpretation of the learning and memory phenotypes. For this reason, we have used a binomial analysis as an alternative method of evaluation.

### Supplemental Figure 2

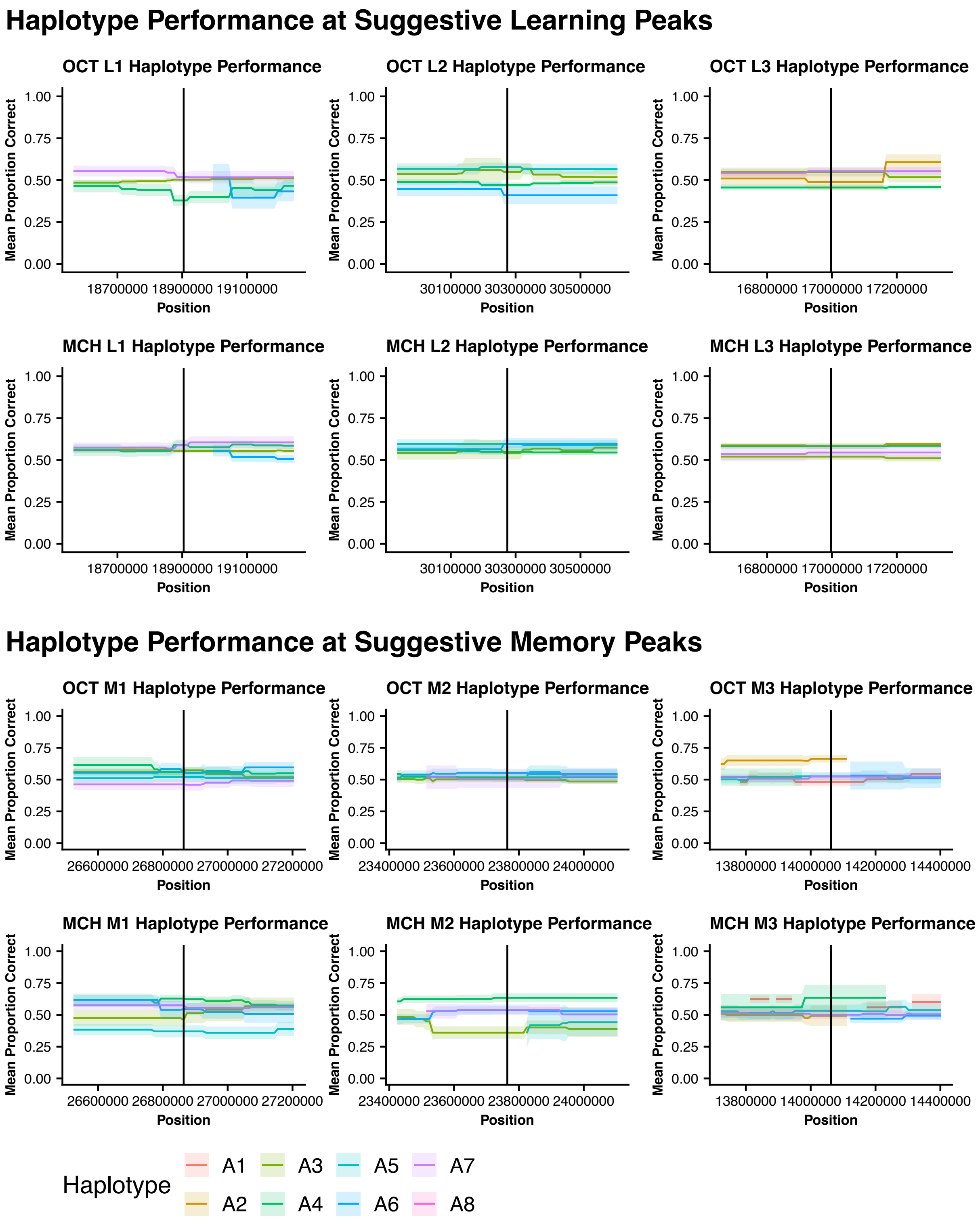


**Supplemental Figure 2.** Haplotype mean performance visualizations for all suggestive olfactory learning and memory peak. Each column compares OCT to MCH performance at 6 identified suggestive peaks.

### Supplemental Figure 3

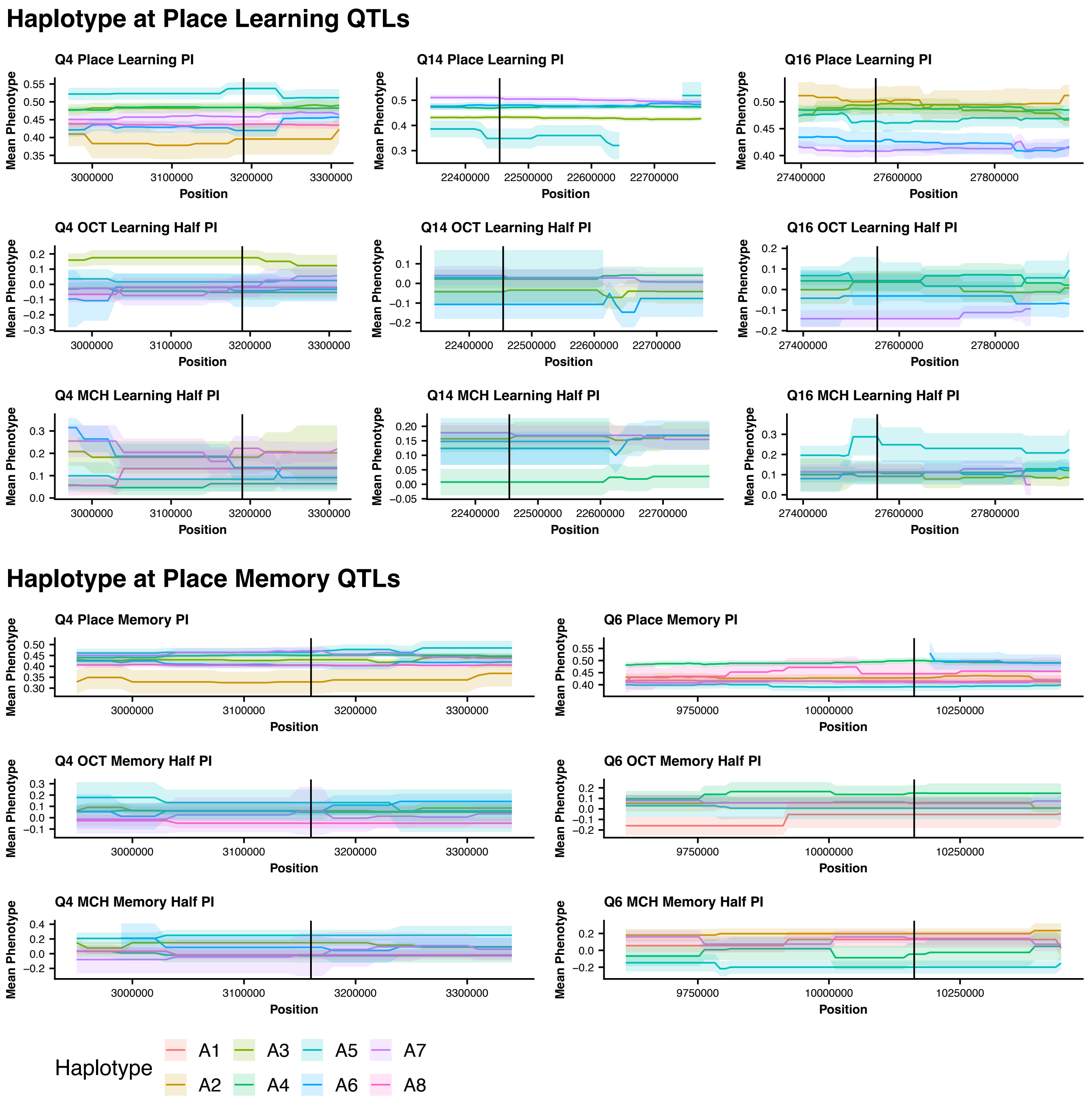


**Supplemental Figure 3.** Haplotype Performance for each phenotype at 4 preselected Place QTLs. Each column compares Place, OCT, and MCH learning or memory phenotype at the preselected QTL positions. Q4 is a shared place learning and memory QTL. Q6 is a place memory specific QTL. Q14 and Q16 are place learning specific QTLs. Important note that scales for place testing and olfactory testing are different.

### Supplemental Figure 4

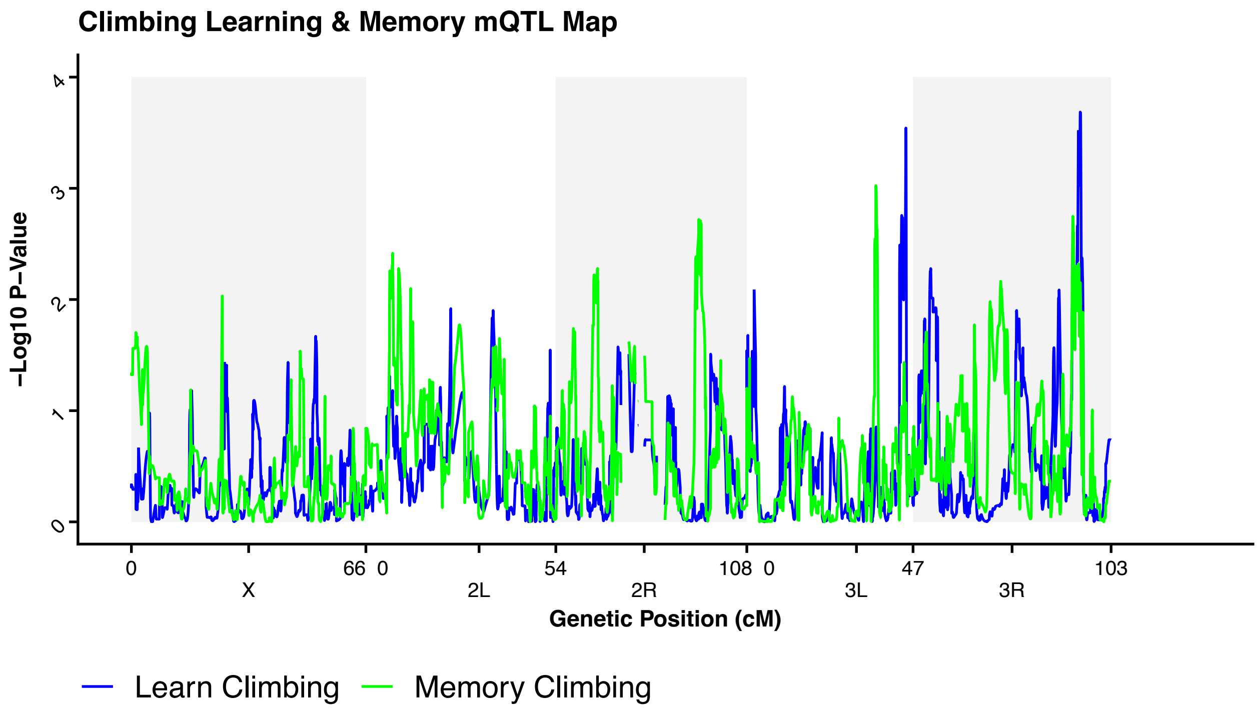


**Supplemental Figure 4.** Climbing Phenotype QTL Map. Climbing during the learning assay is in blue and climbing during the memory assay is green. No significant QTLs were found for climbing.

### Supplemental Figure 5

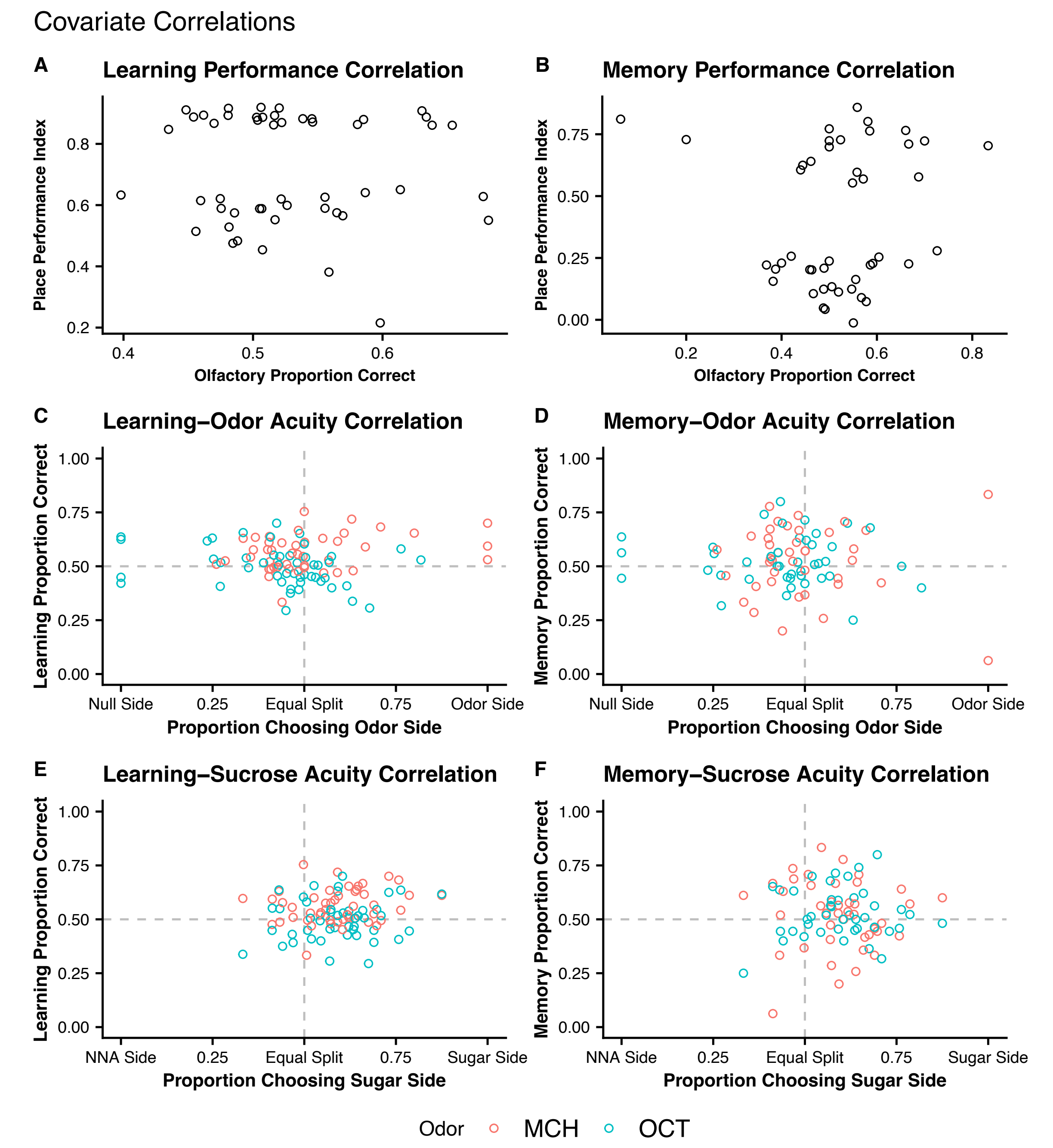


**Supplemental Figure 5.** Performance Correlations. No correlations are found between learning, memory, or acuity tests. **(A)** Place Learning PI scores vs. Olfactory proportion scores. **(B)** Place Memory PI scores vs. Olfactory proportion scores. **(C)** Learning proportion vs. proportion of flies choosing the odor over the null chamber. **(D)** Memory proportion vs. proportion of flies choosing the odor over the null chamber. **(E)** Learning proportion vs. proportion of flies choosing the sucrose sugar agar side. **(F)** Memory proportion vs. proportion of flies choosing the sucrose sugar agar side.
